# Quantifying Drift-Fitness Balance Using an Agent-Based Biofilm Model of Identical Heterotrophs Under Low Nutrient Conditions

**DOI:** 10.1101/2022.12.08.519628

**Authors:** Joseph Earl Weaver

## Abstract

Both deterministic and stochastic forces shape biofilm communities, but the balance between those forces is variable. Quantifying the balance is both desirable and challenging. For example, negative drift selection, a stochastic force, can be thought of as an organism experiencing ‘bad luck’ and manipulating ‘luck’ as a factor in real world systems is difficult. We used an agent-based model to manipulate luck by controlling seed values governing random number generation. We determined which organism among identical competitors experienced the greatest negative drift selection, gave it a deterministic growth advantage, and re-ran the simulation with the same seed. This enabled quantifying the growth advantage required to overcome drift, *e*.*g*., a 50% chance to thrive may require a 10-20% improved growth rate. Further, we found that crowding intensity affected that balance. At moderate spacings, there were wide ranges where neither drift nor growth dominated. Those ranges shrank at extreme spacings; close and loose crowding respectively favoured drift and growth. We explain how these results may partially illuminate two conundrums: the difference between taxa and functional stability in wastewater treatment plans and the difference between equivalent and total community size in neutral community assembly models.

## 1 Introduction

Both stochastic and deterministic assembly processes can shape biofilm populations.^1,2^ Those processes, however, rarely act equally and the balance between them is determined by many conditions related to competition intensity. Such conditions include population size,^3,4^ available space,^5^ and resource availability.^6^ Understanding how this balance shifts under differing conditions provides insights into biofilm-associated systems such as environmental bioreactors, healthcare, industrial production, and natural ecosystems.

Here, we attempt to quantify the balance between drift, a pure stochastic process,^1,3^ and a more deterministic kinetic advantage. Under this balance, even if losing the ‘drift lottery,’ an individual’s progeny may thrive if their maximum growth rate (*μ*_*max*_) or substrate affinity (*K*_*s*_) confers increased fitness over their competitors.

Such quantification is challenging. Drift is an inherently random process and experimental manipulation of a random process, distinct from simply controlling for it, is difficult. Despite that difficulty, there have been some physical experiments in which drift is isolated as an experimental factor,^4,7,8^ often requiring subtle statistical analyses or extremely precise experimental work.

An alternative approach, used here, is to perform the experiments *in silico* where drift may be directly manipulated via random number generation. We used an agent-based model (NUFEB)^9,10^ to simulate spatially competing bacteria under low nutrient conditions. The bacteria were identical and evenly spaced, differentiated only by random growth directions and biomass allocations during division. Drift was therefore the only selection process and was controlled by the seed value initializing the random number generator.

Our goal was to determine the degree to which a deterministic factor (here, Monod kinetics) must improve to overcome negative drift selection, so subsequent simulations using identical seeds were run. The difference was that the ‘biggest loser’, the lineage with the lowest relative abundance, was assigned different kinetics. This approach allowed us to relate quantifiable fitness changes to the likelihood that the failing lineage would overcome negative drift and thrive. We also determined how the required degree of fitness advantage varied under differing crowding intensities (*e*.*g*., closer spacing and increased initial population size).

We found that under purely stochastic conditions the losing lineage varied unpredictably between runs, showing the expected effects of drift. Further, altered fitness did enable losing lineages to overcome drift. For example, for an initial population of 9 cells evenly spaced 10 diameters apart either *K*_*s*_ or *μ*_*max*_ had to improve by at least 10-20% for a 50% chance of thriving. Crowding affected both the improvement needed for a 50% chance of thriving and the ranges over which both drift and fitness co-dominated. The strong and sometimes non-linear interactions between terms could not be adequately reproduced using simple linear estimators but could be adequately expressed with a generalized additive model.

## 2 Methods

### 2.1 Agent Based Model

The agent-based model employed NUFEB (Newcastle University Frontiers in Engineering Biology),^9,10^ which is based on the LAMMPS^9^ molecular dynamics simulation framework and has successfully been used to model multi-species biofilms,^10^ including development and detachment,^7^ trade-offs in extracellular polymeric substance production,^11^ and phototroph-heterotroph metabolic interactions.^12^.

NUFEB is not lattice based, cells were positioned in three dimensions and had individual dynamic sizes. The directions in which cells divided and biomass allocations (40 to 60%) during division were randomly determined using a Park-Miller pseudorandom number generator and were the two factors contributing to drift.

The individually simulated bacterial cells physically interacted using realistic physics and grew according to Monod-style models described by Equation (1) where *μ* is the substrate-dependent growth rate (1/hr), *μ*_*max*_ is the maximum specific growth rate (1/hr), [*S*] is the concentration of the relevant substrate (kg/m^3^), and *K*_*s*_ is the affinity constant for the substrate (kg/m^3^). Additional descriptions of NUFEBs mechanics are detailed in previous publications.^9,10^

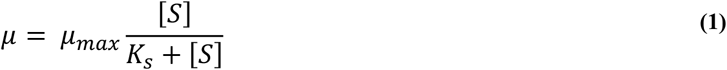

The simulation volume height (2×10^−4^ m) was defined to be in the Z-dimension, the bulk substrate concentration boundary condition at the top of the simulation volume was 1×10^−4^ kg/m^3^ and the initial substrate concentration throughout the volume was set to the same value. The X and Y dimensions were equal and varied based on spacing and number of initial cells. Additionally, the X and Y boundaries were periodic, allowing biomass and substrates to wrap from one side of the simulation to the other.

### 2.2 Experimental Approach

The base experimental unit was an agent-based simulation initially seeded with identical bacterial cells with starting diameters of 1×10^−6^ m, *K*_*s*_ of 3.5×10^−5^ kg/m^3^, *μ*_*max*_ of 1 h^-1^, and yield 0.61 kg biomass per kg substrate consumed. The initial cells (total population 4, 9, or 16) were arranged along evenly spaced (2.5, 5, or 10 cell diameters) *MxM* points at the base of the simulation volume. Bacteria were allowed to grow and compete until 20% of the simulation volume consisted of heterotrophic biomass.

Each combination of populations sizes and spacings was run 120 times using different seed values to initialize the random number generator and the ‘biggest loser’ from each run was identified (see 2.3). Those simulations were then run again, but with the failed lineage given altered kinetic values (see 2.4). The results of the runs were used to determine how the altered kinetics contributed to the probability of transitioning from drift-driven failure to a thriving state (see 2.5) under various crowding intensities.

All combinations of the factor levels listed in **Table 1** (1089 combinations) were simulated for each of the 120 seeds, resulting in a total of 130680 runs. Each run required between 2 to 36 hours to complete, so the simulations were carried out on a high-performance computing cluster (see 2.6).

**Table 1:**
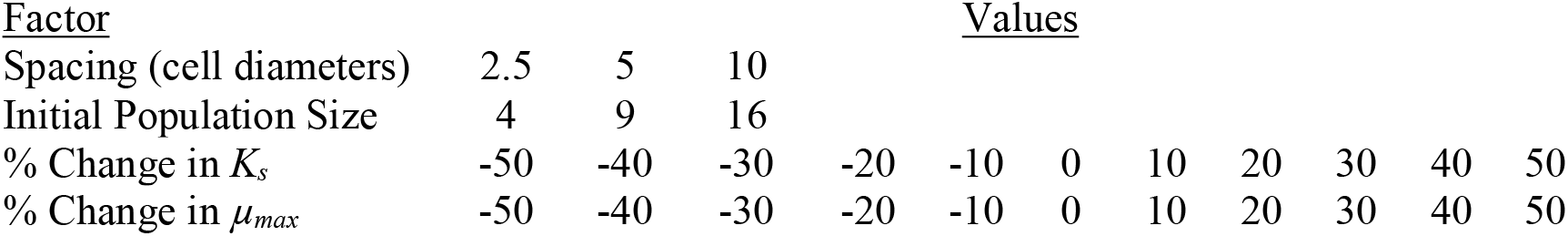
Experimental factors and levels

| <u>Factor</u> | <u>Values</u> |  |  |  |  |  |  |  |  |  |  |
| --- | --- | --- | --- | --- | --- | --- | --- | --- | --- | --- | --- |
| Spacing (cell diameters) | 2.5 | 5 | 10 |  |  |  |  |  |  |  |  |
| Initial Population Size | 4 | 9 | 16 |  |  |  |  |  |  |  |  |
| % Change in $K_s$ | -50 | -40 | -30 | -20 | -10 | 0 | 10 | 20 | 30 | 40 | 50 |
| % Change in $\mu_{max}$ | -50 | -40 | -30 | -20 | -10 | 0 | 10 | 20 | 30 | 40 | 50 |

### 2.3 Determining Failed Lineages

For a system initialized with *N* bacterial lineages, the total biomass *X*_*t*_ is the sum of the biomass for each lineage *X*_*i*_, as expressed by equation(2).

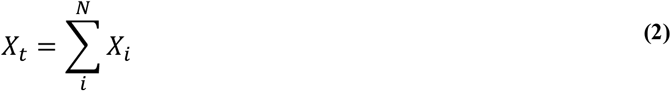

In a system where each initial cell is identical, with no competition, and with no random effects, all *X*_*i*_ are expected to be equal, thus the expected relevant abundance of any lineage (*X*_*E*_) is given as:

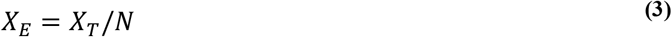

In the first round of simulations, all initial cells were identical and evenly spaced, but cell division directions and biomass allocations during division were determined randomly. As a result, the final biomass for any lineage was often not equal to the expected relevant abundance, *X*_*i*_ ≠ *X*_*E*_. In practice, there were often one or two lineages which strongly dominated with *X*_*i*_ ≫ *X*_*E*_, one or two lineages which became vanishingly small with *X*_*i*_ ≪ *X*_*E*_ (the ‘biggest losers’), and the rest persisted at some noticeable abundance that was however below *X*_*E*._ Moreover, the outcomes appeared to be determined early in the simulation, especially for the best and worst performing lineages. (Supporting Information Figure S1, Table S1, and Video SV1). We have defined three classifications of lineagesurvival based on the difference between *X*_*E*_ and *X*_*i*_: *languishing* (*X*_*i*_ < 0.3 *X*_*E*_), *thriving* (*X*_*i*_ > 0.9*X*_*E*_), and *barely surviving* (0.3 *X*_*E*_ ≤ *X*_*i*_ ≤ 0.9*X*_*E*_).

### 2.4 Fitness Alteration

The worst-performing bacterial lineages from each of the initial homogenous runs were given a potential competitive advantage by altering their individual maximum specific growth rate (*μ*_*max*_) and/or their substrate affinity (*K*_*s*_) (**Figure 1**). The altered values were selected as described in **Table1**. We acknowledge that not all combinations of *μ*_*max*_ and *K*_*s*_ were advantageous and that *μ*_*max*_ and *K*_*s*_ are often strongly correlated; here our goal was to thoroughly explore the parameter space.

**Figure 1:**
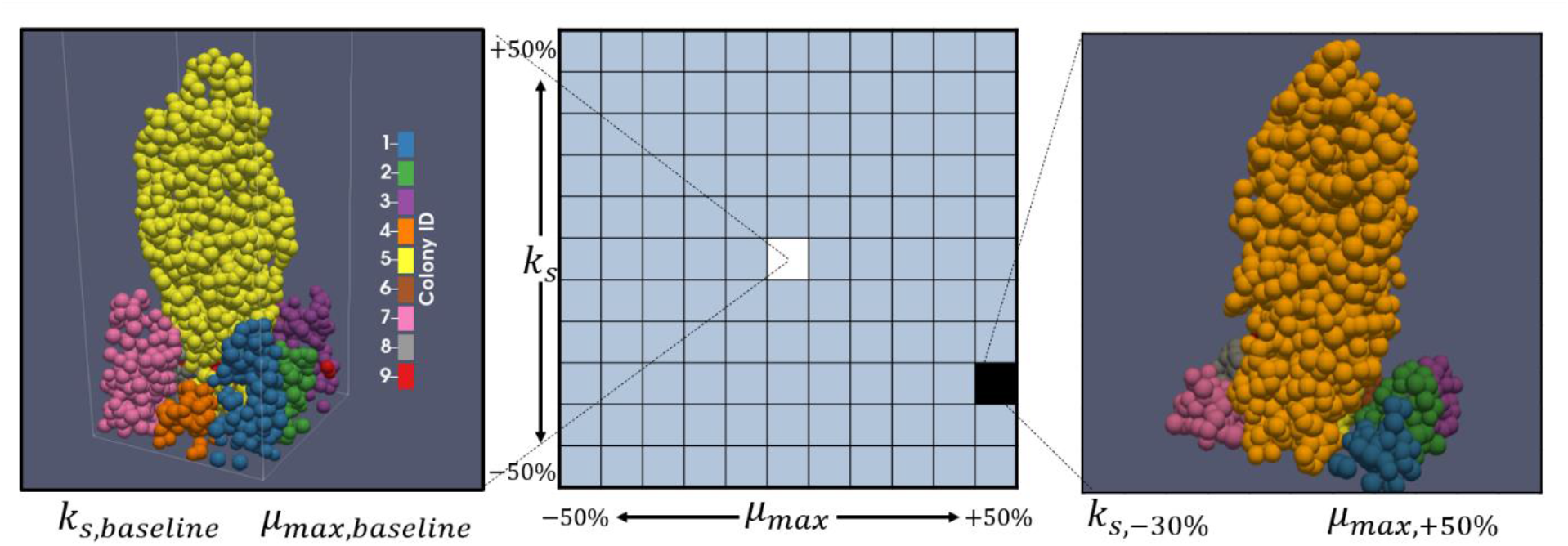
Illustration of a parameter sweep. Under baseline conditions when all bacteria are identical (left hand side), colony 4 was the worst performing lineage. When colony 4 was given a competitive advantage (right hand side) via reduced *KS* and increased *μ*_*max*_, colony 4 transitioned to thriving. This result along with all other parameter combinations across 120 random seeds was used to estimate *p*_*thrive*_, the probability that the worst-performing colony would transition to thriving under given altered kinetics.

### 2.5 Probability Map Generation

The kinetic parameter sweeps were used to generate tables for each combination of factors which listed the final relative biomass of each bacterial lineage, that lineage’s status as the ‘biggest loser’, and the lineage’s success under each run. The percentage of failing lineages across all random seeds which transitioned to thriving was calculated for each combination of initial population size, spacing, *μ*_*max*,_ and *K*_*s*._ Those percentages represent the probabilities that the fitness advantage (if any) conferred by altered kinetics would outweigh negative drift selection under the given conditions.

### 2.6 Simulation Management

Simulations were run and their results tabulated on the Newcastle University Rocket High Performance Computing environment and managed using Snakemake^13,14^ workflows populating a SLURM^15^ queue. Each simulation was run on a single core, with multiple hundreds of simulations run in parallel. Job submissions encompassed all kinetic parameter sweeps for each combination of other parameters, *e*.*g*., a single batch submission would consist of all combinations of *μ*_*max*_ and *K*_*s*_ for 4 bacteria, spaced 5 diameters apart.

### 2.7 Data Analysis

Simulation results were saved as tabular comma separated value (CSV) text files and aggregated using BASH^16^ (v. 4.2) shell and Python^17^ (v. 3.8) scripts which included the NumPy^18^ and pandas^19^ libraries. Further processing of the data was performed off the cluster and used R^20^ (v. 4.2) scripts incorporating various Tidyverse^21^ and other supporting packages.^22–43^

#### 2.7.1 Parameters Quantifying the Balance Between Drift and Fitness

Each probability map was conceptually analogous to a cliffside; a continuous sharp probability threshold gradient separated by two flat regions of either 100% lineage success or failure (**Figure 2** A). We wished to quantify the midpoint and steepness of the gradient along lines of constant *K*_*s*_ for each crowding condition. A cross-section of the probabilities along *μ*_*max*_ for any constant *K*_*s*_ produces a sigmoid-shaped profile (**Figure 2** B). The profiles were fit to a logistic function of *μ*_*max*_ with a maximum value of 1 given by equation (**4**), where *p*_*thrive*_ is the probability of transitioning to a thriving colony, *k* is a parameter affecting the steepness of the curve, and *μ*_*50*_ is the *μ*_*max*_ value at which there is a 50% probability of thriving.

**Figure 2:**
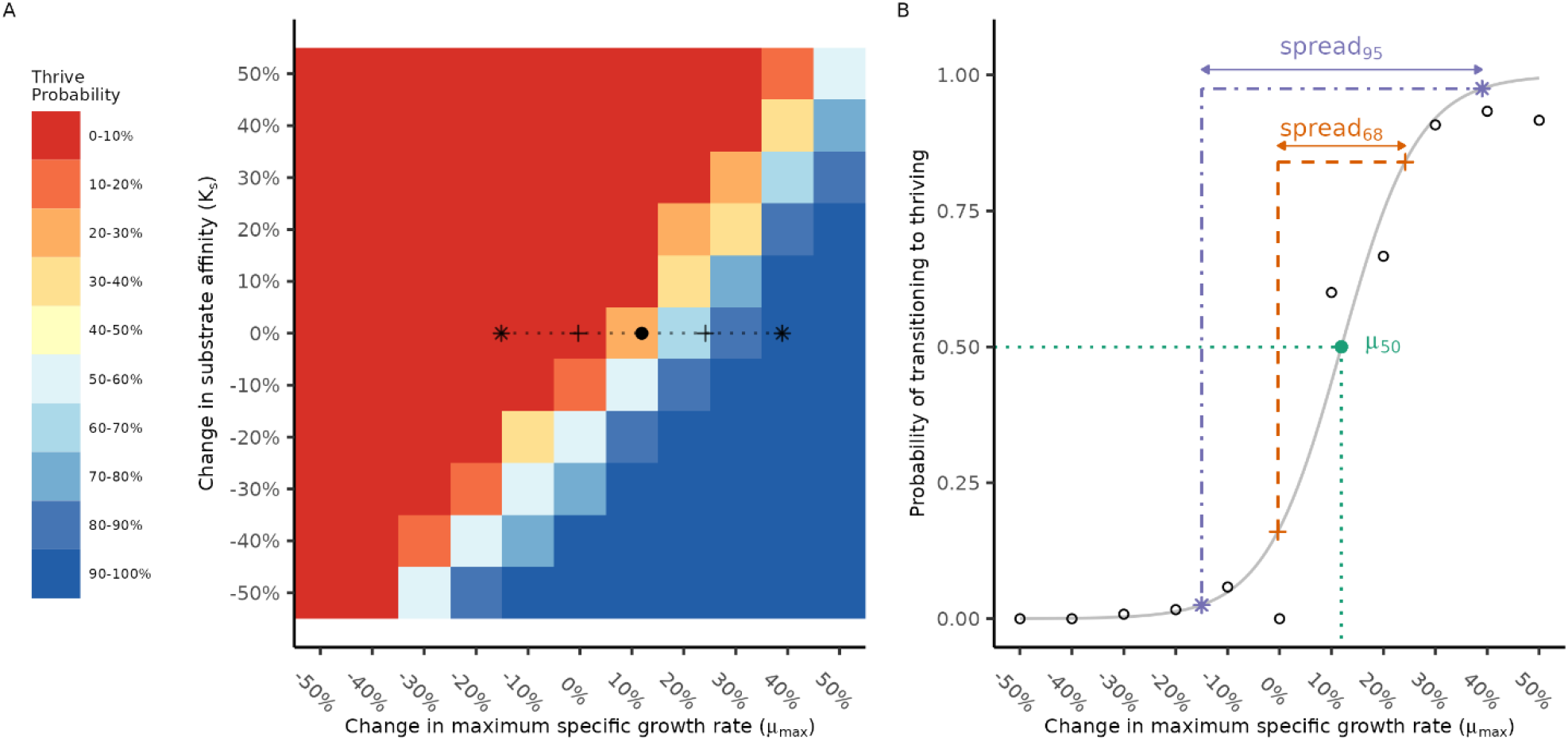
Illustration of how the *μ*_*50*_ and *spread* parameters were calculated. In this example, the probability map corresponding to 4 initial organisms placed 5 diameters apart is shown (A), and the dashed line is drawn along a line of constant *K*_*s*_. The full length of the line denotes the *spread*_*95*_ region, the portion between crosses denotes *spread*_*68*_, and the solid point represents the *μ*_*50*_ mark. When the *p*_*thrive*_ values are plotted as a function of *μ*_*max*_ along the line of constant *K*_*s*_, (B) it is apparent that a logistic function (grey solid line) may be fitted to the points (black rings). The fitted function was used to estimate both the value of *μ* corresponding to *μ*_*50*_ and the widths of the *spread* regions. This analysis was repeated for all crowding conditions along all lines of constant *K*_*s*_.

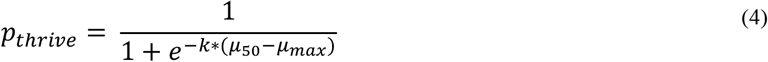

The relevant *k* and *μ*_*50*_ parameters from each fit were recorded. We also determined the domains of *μ*_*max*_ values associated with the *p*_*thrive*_ ranges covering either a 2.5-95% or 16-84% chance of thriving. These domains, respectively named *spread*_*95*_ and *spread*_*68*_ quantified the regions over which neither drift nor fitness dominated.

The results of all sigmoid fits are shown in Supporting Information Figures S2-S10.

#### 2.7.2 Analysing Balance Parameters

Within each crowding scenario, the extracted parameters were analysed using simple linear regression models of the parameters as functions of *K*_*s*_. The effect of crowding pressure (spacing and total population) was then analysed by comparing the results of the fits between scenarios.

We note that although the linear fits for a 2^nd^ order polynomial on *μ*_*50*_ generally resulted in marginally improved *R*^2^ scores and removed parabolic patterns from the residuals, the simple linear regressions were still excellent and more interpretable; care should be taken if extending this work to larger ranges of kinetic values.

#### 2.7.3 Modelling the Effect of Competitive Pressure and Altered Kinetics

We wished to determine if a model based on the simulation results could accurately reproduce the transition probabilities for each crowding scenario. The ultimate goal of these models was not prediction, but to provide a descriptive framework^44^ showing which factors, interactions, and potential non-linearities were important. Variations on both multiple linear regression models (MLR) and Generalized Additive Models (GAMs)^45^ were fitted to either the log-likelihood of *p*_*thrive*_ (for MLRs) or directly to *p*_*thrive*_ (GAMs).

In both cases, backward step selection from factorial models incorporating up to three-way interactions was performed to select the final model. Non-significant (p > 0.05) terms were iteratively removed from the model starting with the highest order interactions. Main effects were retained even if non-significant when they were part of a significant interaction term.

The final models were selected based on *R*^2^ and Akaike Information Criterion (AIC) values as well as interpretability. The potential models and the associated fit criteria are included in Supporting Information Tables S2-S5.

## 3 Results

### 3.1 Drift Occurred When All Cells Were Identical

A foundational assumption of this approach is that even in a system with equally spaced, identical microbes, random growth will lead to drift. We tested this assumption for crowding scenarios where all microbes had identical base *K*_*s*_ and *μ*_*max*_ parameters by determining the number of times each lineage was the ‘biggest loser’ over 120 simulations (**Figure 3**) and, similar to testing *m* dice for fairness, applied a Chi-Square test (α=0.05/*m*) where *m* is a Bonferroni correction for multiple testing (*m*=9 at 3×3 initial spacings and population sizes). Each initial site was statistically as likely as any other to be the biggest loser (Supporting Information Table S6).

**Figure 3:**
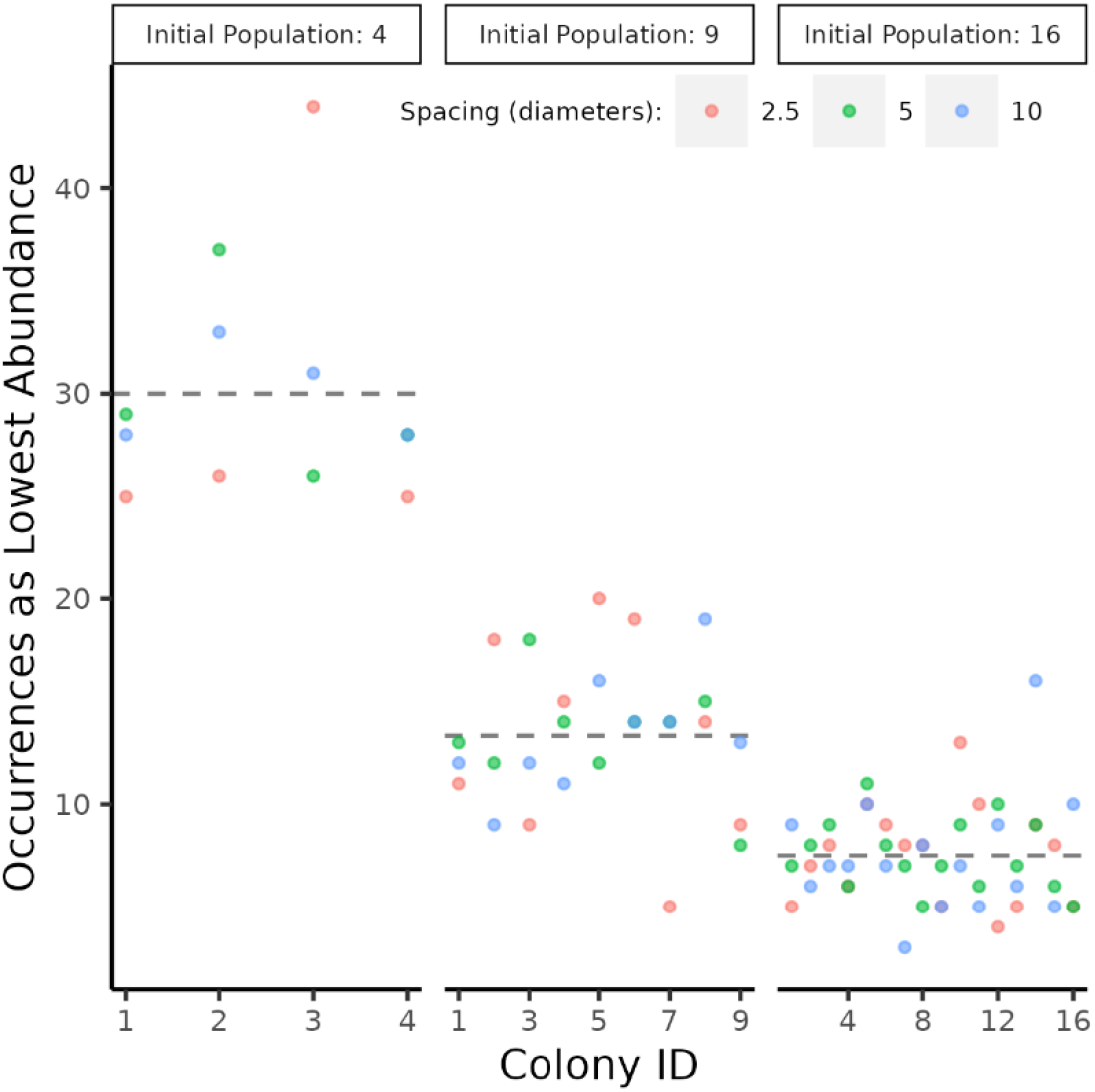
The number of times each colony was the least successful performer during all 120 runs of the baseline simulation where all bacteria were identical. Dashed grey lines indicate the expected value. Points are colored based on spacings between initial sites. For each set of initial populations, no colony appeared biased away from the expected number of failures.

Additionally, the relative proportion of lineages which languished, survived, or thrived for each set of crowding conditions was determined. Simulations, on average, had between one and two thriving lineages, with the rest languishing (65-75% for 4 initial sites, 80-88% others), and a few (0-5%) which did not thrive but grew to non-negligible abundance (Supporting Information Table S1). When 4 organisms were initially present, only languishing and thriving lineages existed, there was otherwise no clear trend between these ratios and either the number or spacing of initial bacteria.

### 3.2 The Least Successful Lineages Could Overcome Drift with Altered Kinetics

As expected, improving the relative fitness of an organism gave it a chance to overcome negative selection via drift (**Figure 4**).

**Figure 4:**
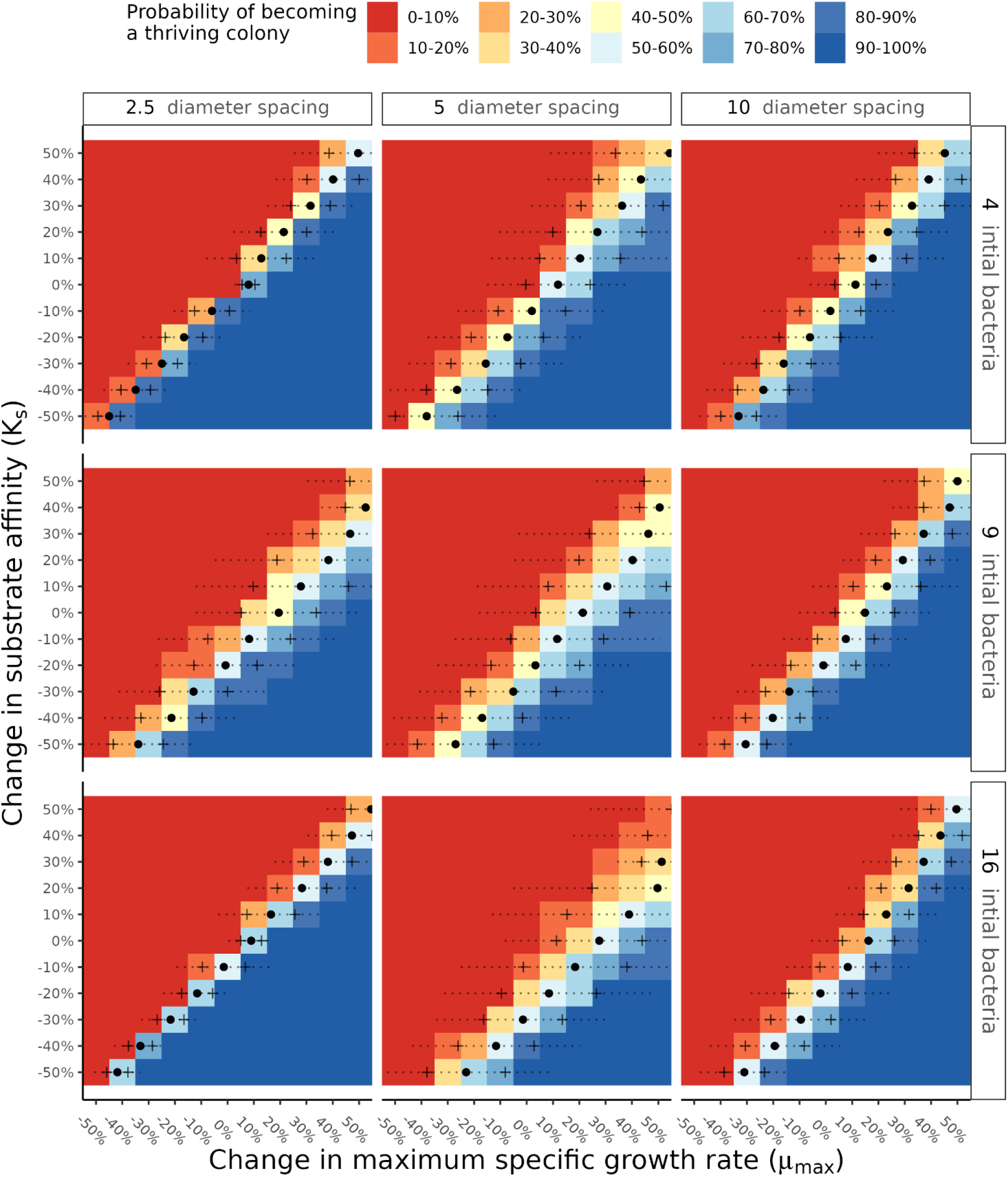
Changing the *μ*_*max*_ and *K*_*s*_ of the least successful lineage was associated with a probability of transitioning to a thriving status. Solid dots represent *μ*_*50*_, the percent change in *μ*_*max*_ at a given *K*_*s*_ associated with 50-50 odds of thriving. Dashed lines show the range of *μmax* corresponding to a *p*_*thrive*_ of 2.5 to 97.5 (*i*.*e*., *spread*_*95*_). Crosses indicate the analagous *spread*_*68*_ region.

The increases in *μ*_*max*_ corresponding to the least successful lineage having a 50% chance to become thriving, which we denote as *μ*_*50*_, are represented by the dark circles in **Figure 4**. At the baseline *K*_*s*_ a typical *μ*_*50*_ is in the range of 10-30%, with the exact value affected by initial spacing and population size (*i*.*e*., crowding). Decreasing *K*_*s*,_ as expected, reduces *μ*_*50*_ – even to the point where so long as substrate uptake affinities are ‘good enough’, the initially failing organism may have excellent odds despite having a *μ*_*max*_ notably lower than its peers. The overall effect, for a given crowding condition, is a semi-linear ‘cliff ‘of *μ*_*50*_ values where *μ*_*50*_ changes inversely with *K*_*s*_. Qualitatively speaking, the location of that ‘cliff’ was shifted to the right (higher *μ*_*50*_) when crowding was increased either via initial population size.

Areas where the probability of thriving is neither 0 (drift dominated) nor 1 (fitness dominated), are, by definition, areas where drift and fitness both determine success. The widths of these areas are denoted as *spread* and are indicated by the dotted horizontal lines and crosses in **Figure 4**. The full length of the line denotes the *spread*_*95*_ area, which is the range of *μ*_*max*_ for a given *K*_*S*_ which corresponds to a 2.5% to 97.5% chance of thriving. The crosses represent a similar range, *spread*_*68*_, which corresponds to a 16% to 85% chance of thriving.

Because the *μ*_*50*_ values are also the centre point of the *spread* regions, spread shifted in the same manner as *μ*_*50*_. However, the actual magnitudes of *spread* did not necessarily follow the same patterns. First, there was no guaranteed symmetry about *K*_*s*._ For example, for 9 initial organisms separated by 5 diameters, the *spread*_*95*_ for *K*_*s*_ of -30% and 30% are visibly different (**Figure 4**, row 2 column 2). Though the asymmetry varied between crowding conditions, it generally manifested as spread widening with increasing *K*_*s*_. Second, there was no clear monotonic trend with spread values corresponding to crowding. A spacing of 5 diameters appeared to produce the widest spreads, *ceteris paribus*. Further, there was no clear rule determining which of the two spacing extremes would have a larger *spread*. For example, with 4 initial bacteria a spacing of 10 diameters resulted in larger spreads than in 2.5 diameters, but the opposite occurred with 16 initial bacteria.

### 3.3 Quantitative Effect of Crowding on *μ*_*50*_ and s*pread*

The qualitative effects of crowding described in the previous section were quantified via simple linear regression as described in section 2.7.2.

For any given crowding condition *μ*_*50*_, the relative change of *μ*_*max*_ at which the worst performing lineage had a 50% chance to transition towards thriving, was essentially linear with respect to *K*_*S*_ and the correlation coefficient was uniformly high (**Figure 5**). The slopes of these relationships indicate the change in *μ*_*50*_ required to compensate for a change in *Ks*. At the tightest spacing, *μ*_*50*_ had to change the most, with a ratio of essentially 1:1 and a slight monotonic increase corresponding to initial population size. As initial spacings widened, the ratio almost always decreased for any initial population size. Across initial population sizes, the ratio for 5 and 10 diameter spacings appeared to follow a general trend of increasing, but this was not monotonic.

**Figure 5:**
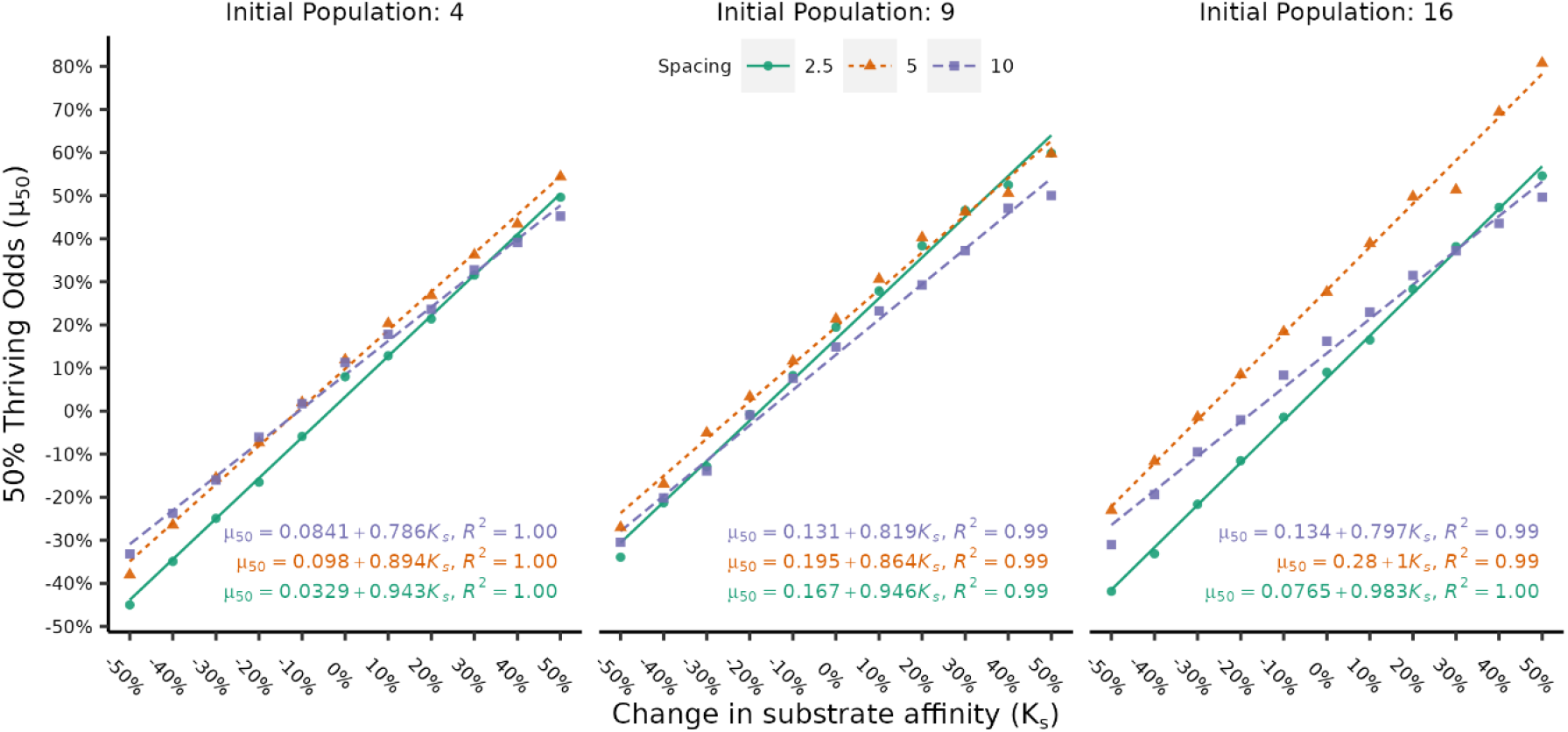
Under each crowding condition, *μ*_*50*_ changed linearly with *K*_*S*_. Large initial population sizes increased the differences between spacings, moderate spacings generally required the largest absolute *μ*_*50*_, but the tightest spacings required the largest change *μ*_*50*_ in per unit change in *K*_*S*._

The absolute value of *μ*_*50*_ was strongly affected by differences between the fitted intercepts. For example, a 2.5 diameter spacing under an initial population size of 16 had a high slope (0.983) but also the lowest required *μ*_*50*_ of all spacings under the same conditions until a 30% change in *K*_*s*_. The practical difference between spacing was largest at high initial population size, indicating a potential interaction between these factors.

Unlike *μ*_*50*_, the range over which both drift and fitness effects co-dominated, *spread*_*95*_ did not have a simple linear relationship with *K*_*S*,_ with many poor *R*^2^ values, residual patterns, and high leverage datapoints (**Figure 6**). There was also no clear, consistent relationship applicable across factors. In general, linear fits became worse with increasing population size which appeared to produce higher variance and generated more high-leverage points, especially at separation distances of 5 diameters. These issues were largely the same when the analysis was repeated for *spread*_*68*_ (Supporting Information Figure S13). There is little to concretely say except that the *spread* was most often widest at moderate spacings, generally increased with *K*_*s*,_ and had a noisy, complicated relationship with initial population size and spacing.

**Figure 6:**
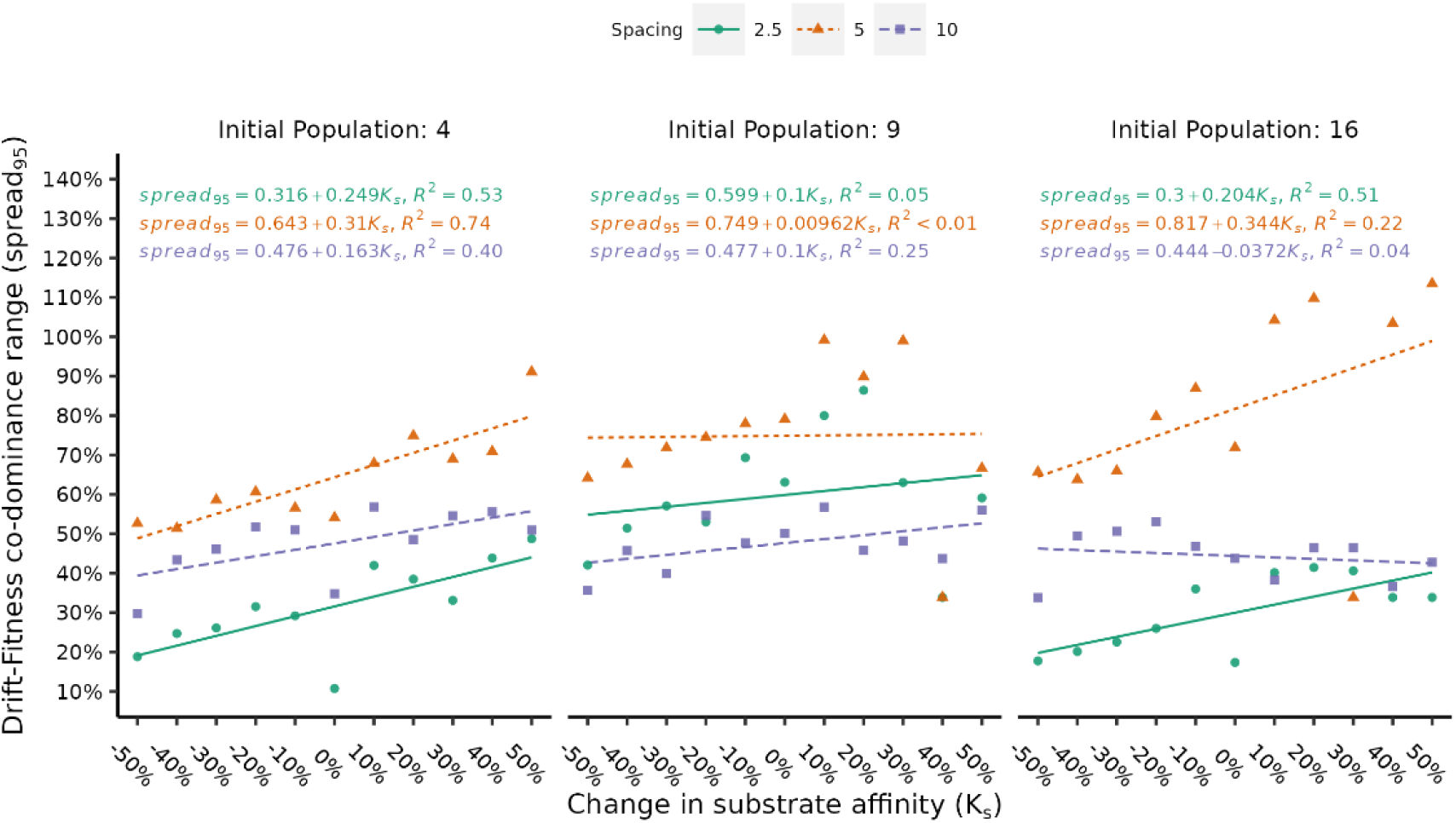
Under each crowding condition, *spread*_*95*_ changed with *K*_*S*_. Insofar as trends were present, moderate spacing produced the widest *spread*_*95*_ and the differences between spacings increased with population size.

### 3.4 Description via Multiple Linear Regression and Generalized Additive Models

The simulation results were modelled using both multiple linear regression (MLR) and a generalized additive model(GAM) respectively described by equations (**5**) and (6) where: *p*_*thrive*_ is the probability of transitioning to a thriving status, *μ*_*p*_ and *K*_*p*_ are the respective percent changes from the baseline *μ*_*max*_ and *K*_*s*_, *N*_*0*_ is the initial population size, *s*_*i*_ is the initial spacing (in diameters) between organisms, and ε is a small pseudo-probability (1×10^−6^) added to avoid division by 0 and issues with log transformation. For linear terms in equations (**5**) and (6), *β*_*i*_ denotes the fitted coefficient for term *i* with *i*=0 representing the intercept. Terms to which GAM smoothing was applied are represented by s(*…*) in equation (6) with interactions between a smoothed variable *x* and linear variable *y* denoted as s*(x*, by *y)*. Significant terms (p < 0.05) are highlighted in bold. The associated coefficients, significance values, and other relevant fitting information are included in Supporting Information Tables S2-S5.

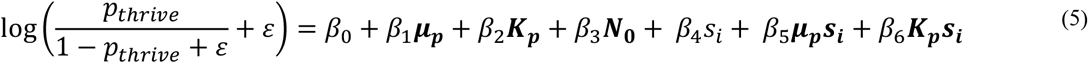

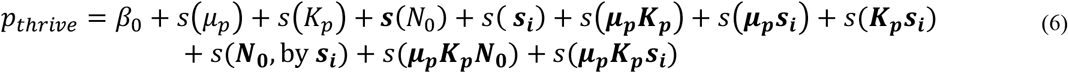

The MLR model captured the general behaviour of the shift in the boundary between low and high thriving probabilities but did not adequately reproduce changes in *spread* (**Figure 7** A *vs*. C). The overall root-mean-squared error (RMSE) of the model was 0.125. While most predicted probabilities differed from the simulation by no more than ±0.1, some predictions were subject to large error (**Figure 7** A, D, F and Supporting Information Figures S11 and S14-S15). The largest errors unsurprisingly appear closest to the boundary between low and high *p*_*thrive*_ regions with the MLR model over-optimistic at the extremes of spacing and lower initial population size. Conversely, the model tended towards overly pessimistic at moderate spacing.

**Figure 7:**
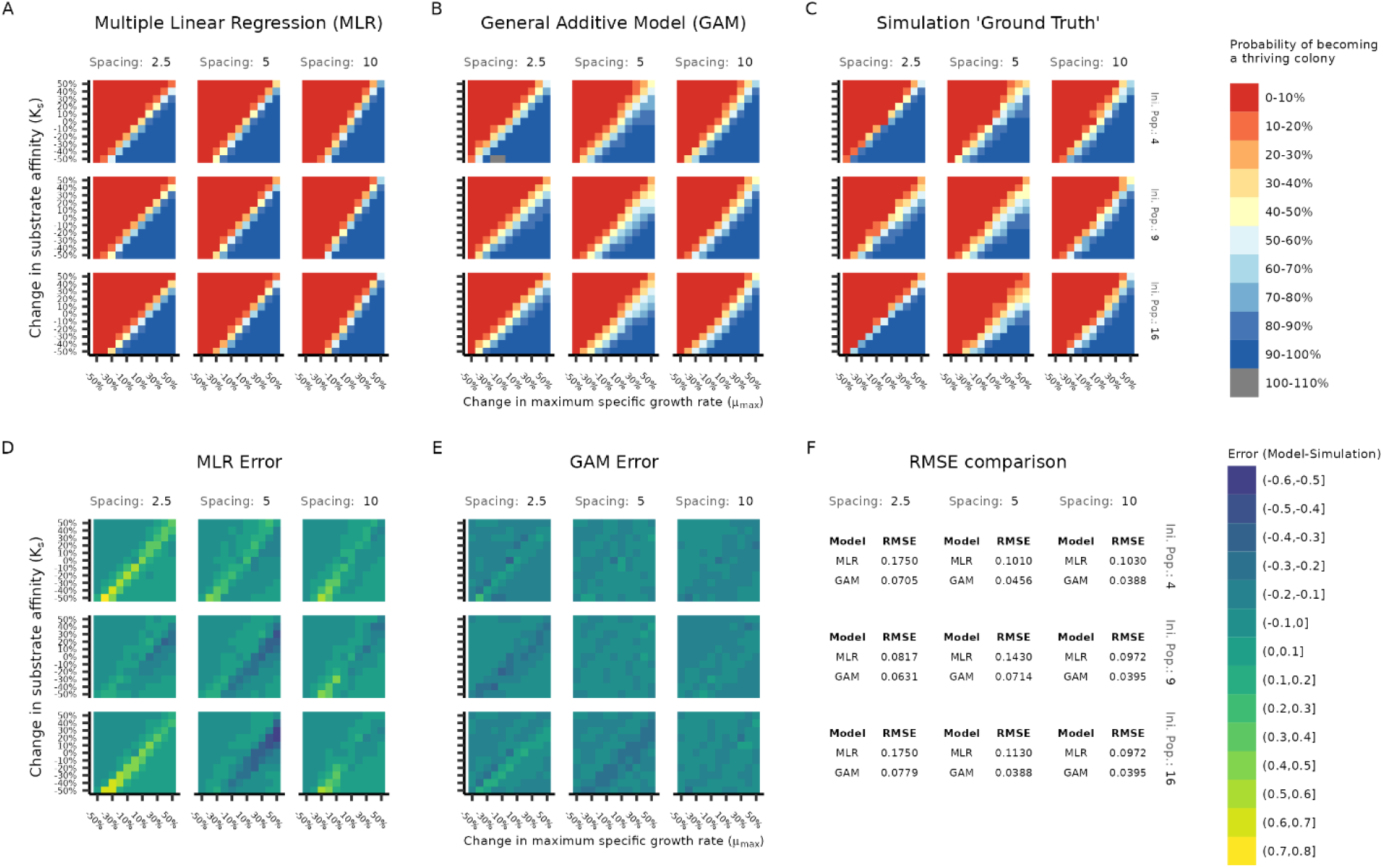
Predictions of MLR model (A) and GAM (B). Simulation results in (C) are presented for ease of comparison. The model errors for the MLR (D) and GAM (E) are presented visually as well as quantified per-crowding condition in (F). The GAM outperformed the MLR, which particularly failed to capture *spread*, was overly optimistic at spacing extremes, and pessimistic at moderate spacing. The small region of greater than 100% odds occured because the GAM was not constrained to predicting values in the range of [0,1]. Larger individual plots of panels A, B, D, and E are available in Supporting Information figures S14-S17.

In comparison to the MLR model, the GAM not only captured the general boundary shift but also the changes in *spread* (**Figure 7** B *vs*. C in contrast to A *vs*. C). The overall RMSE of the GAM was 0.0563, or somewhat better than half the RMSE of the MLR model. As with the MLR model, most predicted probabilities differed from the simulation by no more than ±0.1. Unlike the MLR model, there were fewer exceptionally large errors and those which did occur were of smaller magnitude (**Figure 7 B**, E, F and Figures S12 and S16-S17). The GAM followed the same trends in over- and under-prediction as the MLR.

## 4 Discussion

### 4.1 Crowding Affects the Balance Between Drift and Need for Fitness

The two parameters describing the balance between drift and fitness, *μ*_*50*_ and *spread*, were both affected as crowding became more intense due to either decreased initial spacing or increased initial population size. It was originally expected that as crowding intensity increased, greater fitness would be required (*μ*_*50*_) along with a decrease in the range of values over which both drift and fitness co-dominate (*spread*). That was not the case.

Instead, the largest *spread* values predominately occurred at moderate (5 diameter) initial spacing. We suggest the cause is physical competition for space, specifically the practical significance of single ‘bad’ random choices in division direction and biomass allocation. When bunched tightly together, competition for space is intense and even a few poor random events can consign a lineage to languishing despite a moderate growth advantage. At the other extreme, spatial competition is lessened sufficiently that a few missteps do not guarantee ruin, allowing a lineage to take the full benefit of any growth advantage. Meanwhile, at moderate spacing, immediate neighbours are close enough so that poor random events are harmful but not necessarily disastrous and, at the same time, growth advantages are somewhat hindered, but still helpful. Remembering that spread quantifies the region where neither fitness nor drift dominate, it then makes sense that we observed the largest spread values at moderate spacing.

The 50-50 odds point, *μ*_*50*_, was also slightly larger at moderate spacings, although not consistently and the effect size was not practically different except at large population sizes. The underlying basis for why is not entirely clear, numerically it was due to the consistently larger intercept (Figure 5). The trend of the slopes is, however, more easily explained and we attribute it to competition for substrate. For any initial population size, smaller spacings resulted in higher slopes. In other words, to maintain the 50-50 odds when *K*_*s*_ was poor, *μ*_*50*_ had to change more at closer spacing. This makes intuitive sense – closer spacings result in lower local substrate concentrations, and any deficit to *K*_*s*_ is more deleterious to fitness.

Increased initial population sizes had more straightforward, secondary, effects on *μ*_*50*_ and *K*_*s*_. As the initial population size increased, the differences between spacings became more pronounced, but the general trends remained unchanged. In other words, more competitors are problematic, especially as it relates to diffusible substrate, but the major influence on success is competition for space between immediate neighbours.

### 4.2 Interactions Between Factors Incorporating Non-Linear Effects are Important

In the MLR a main-effects only model (RMSE 0.125, *R*^2^ of 0.820) performed essentially identically to the MLR model with interactions (RMSE 0.127, and *R*^2^ of 0.820), however neither adequately reproduced simulation results and were especially poor at representing the regions where both fitness and drift co-dominated. A GAM which incorporated only main effects using non-linear smoothing quantitatively performed slightly worse than either MLR main-effects model (RMSE 0.197 and *R*^2^ of 78.1), but drastically and uniformly overpredicted spread. Only when both interactions and smoothing were incorporated did a model adequately reproduce the simulation results (**Figure 7** and Supporting Information Figure S17). It is visually apparent in the simulation results and quantified in the fitting results (Supporting Information Table S4-5) that interactions are important, particularly those involving spacing. Further, the non-linearity of the interactions (measured as the departure of the term’s extended degrees of freedom from a value of 1), is particularly high for any interaction incorporating both *μ*_*p*_ and *K*_*p*_ and less so but still notably for interactions incorporating spacing (Supporting Information Table S5).

### 4.3 Limitations and Extensions

The simulated conditions were deliberately chosen to isolate the effect of drift. While this made the work tractable, a system wherein every organism is completely identical, starts growing at the same time, and is initially evenly spaced on a grid does not frequently occur in nature. Although we believe the general themes uncovered translate to real ecological systems, the exact quantification does not and is not mean to apply to all situations. Future work should focus on stochastically placed (in time and space) populations with natural variability in Monod parameters.

Extending the work so that the simulated community reflects a more natural distribution would also enable validation of the model, as, despite promising advances,^46^ it is currently infeasible to exactly place essentially identical bacteria at the resolution required.

Additional parameters affecting drift and fitness should also be evaluated – especially the influence of nutrient-rich conditions^47^ and how a change to yield, rather than growth rate, alters success.^48^ Adding these factors requires however overcoming the curse of dimensionality, the current simulations took over 1 year of real-world time and 175 years’ worth of CPU time. Given the large areas where ‘nothing interesting’ happens, designing further experiments to incorporate adaptive sampling^49^ is a promising solution. Further, adaptive sampling would enable, at the same computational cost, exploring a larger range of *μ*_*max*_ and *K*_*S*_ variation (which may vary by orders of magnitude in real-world conditions^50^) and at a greater degree of resolution than 10% changes in the region where the probabilities rapidly change.

## 5 Conclusion And Relevance to Real World Systems

It is apparent that during biofilm formation in low nutrient conditions, drift strongly determines which organisms thrive and which organisms fail, so long as they have similar growth rates and substrate affinities. Even when those parameters differ between individuals by ±50%, there are still large regions where a fitness advantage does not guarantee overcoming negative drift selection.

In fact, we observed the lineage fates were determined very early in the simulations and for these systems ‘well-begun is half done’. We speculate that this may be a piece to the puzzle explaining the apparent contradiction between actual and effective community size in neutral modelling^4^ – the bacteria are not in competition with the full steady-state community but only the immediate smaller, community near the beginning of biofilm growth. However, the conditions studied here violate the steady state assumption of that work, so a more careful analysis is warranted.

The conditions we have described are not dissimilar from those within an aerated portion of a wastewater treatment plant, where tightly packed bacterial aggregates are suspended in a bulk liquid and where substrate concentrations are often quite low, especially during operation as a completely mixed stirred reactor (albeit somewhat higher than simulated here). Further, these bacteria are recirculated through the system and relatively well-adapted to domestic wastewater, thus already selected for similarity. Based on the results presented here, we would expect to see a system in which there is a high degree of random turnover in organism identity, but relatively stable functional and biological activity, which is exactly what has been observed in wastewater treatment plants.^51,52^

## 6 Acknowledgements

We wish to acknowledge the United States National Science Foundation Directorate for Biological Sciences for funding via the Postdoctoral Research Fellowships in Biology (NSF PRFB Award # 2007151) and the Newcastle University Rocket HPC computing cluster. We also thank Tom Curtis for his valuable feedback and encouragement, Denis Taniguchi and Bowen Li for their help in understanding the NUFEB software, and countless helpful comments from many interested parties during poster sessions and talks.

## 7 Competing Interests

The author has no competing interests.

## 8 Data Availability Statement

The data analysis code, data from the simulations, and exact NUFEB variant are respectively located in the following repositories:

Analysis: https://github.com/joeweaver/agent_based_biofilm_drift/ Data: https://osf.io/fch3z/

NUFEB variant: https://github.com/nufeb/NUFEB-dev/tree/compute_vol_group

## Notes

https://osf.io/fch3z/

https://github.com/joeweaver/agent_based_biofilm_drift/

https://github.com/nufeb/NUFEB-dev/tree/compute_vol_group

### Competing Interest Statement

The authors have declared no competing interest.

