## Supporting Information for "Quantifying Drift-Fitness Balance Using an Agent-Based Biofilm Model of Identical Heterotrophs Under Low Nutrient Conditions"

Growth Example Over Time

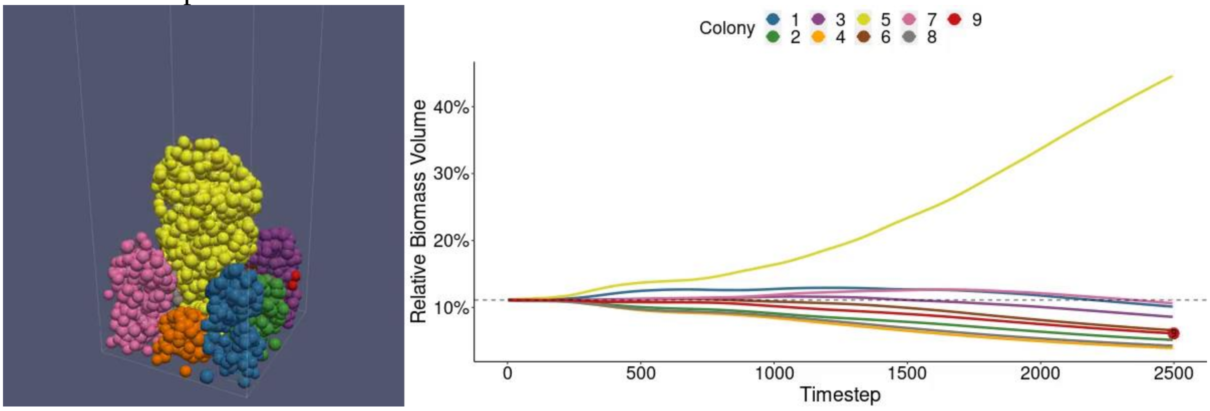


**Figure S1:** Example of growth over time in a simulation. Here a 3x3 grid of identical heterotrophs (seed 1701) was grown and the relative biomass over time was tracked. Dashed grey line indicates the expected relative biomass if evenly distributed between lineages. An animated version is available as a supporting video, SV1.

#### Ultimate Fates of Organisms

**Table S1:** Percentages (mean ± std. deviation across runs) of bacterial lineages which thrived, languished, or barely survived for each combination of initial total population and spacing.

| Initial Population | % Thriving Lineages (by Spacing) | | | % Languishing Lineages (by Spacing) | | | % Barley Surviving Lineages (by Spacing) | | |
|---|---|---|---|---|---|---|---|---|---|
| | 2.5 | 5 | 10 | 2.5 | 5 | 10 | 2.5 | 5 | 10 |
| 4 | 35.2 ± 13.2 | 26.7 ± 6.3 | 25 ± 0 | 64.8 ± 13.2 | 73.3 ± 6.3 | 75 ± 0 | 0 ± 0 | 0 ± 0 | 0 ± 0 |
| 9 | 11.9 ± 2.8 | 11.9 ± 2.8 | 14.9 ± 5.7 | 84.3 ± 6 | 84.3 ± 6.2 | 81.5 ± 7.9 | 3.9 ± 5.3 | 3.9 ± 5.3 | 3.6 ± 5.2 |
| 16 | 14.8 ± 6.6 | 7.7 ± 2.7 | 13.8 ± 5.2 | 81.3 ± 7.9 | 88.4 ± 4.1 | 80.3 ± 7.9 | 3.9 ± 3.5 | 3.9 ± 3 | 5.9 ± 4.2 |

#### Sigmoid Fits

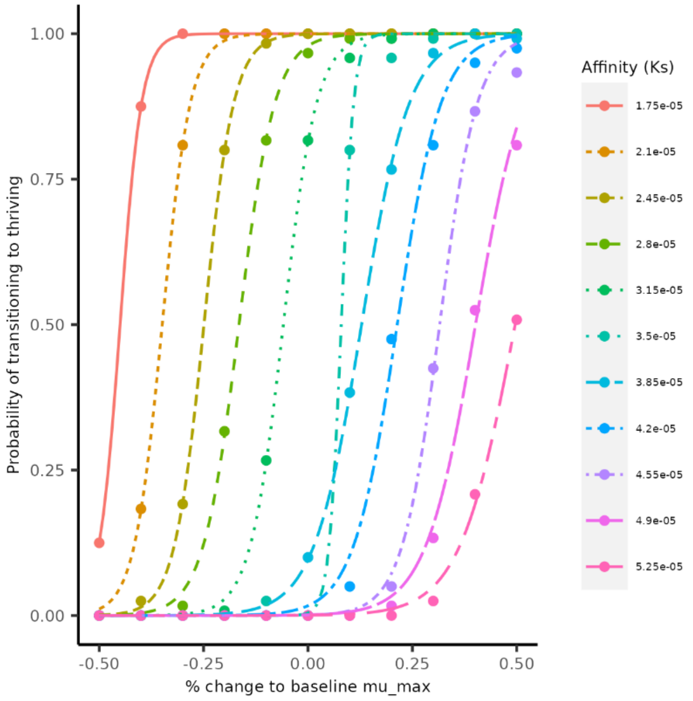


**Figure S2:** Sigmoid fits for 4 initial organisms spaced 2.5 diameters apart.

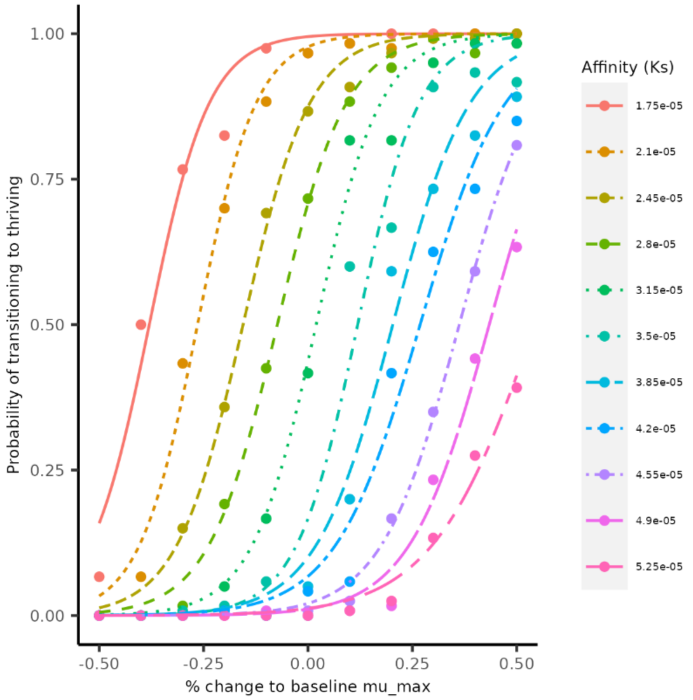


**Figure S3:** Sigmoid fits for 4 initial organisms spaced 5 diameters apart.

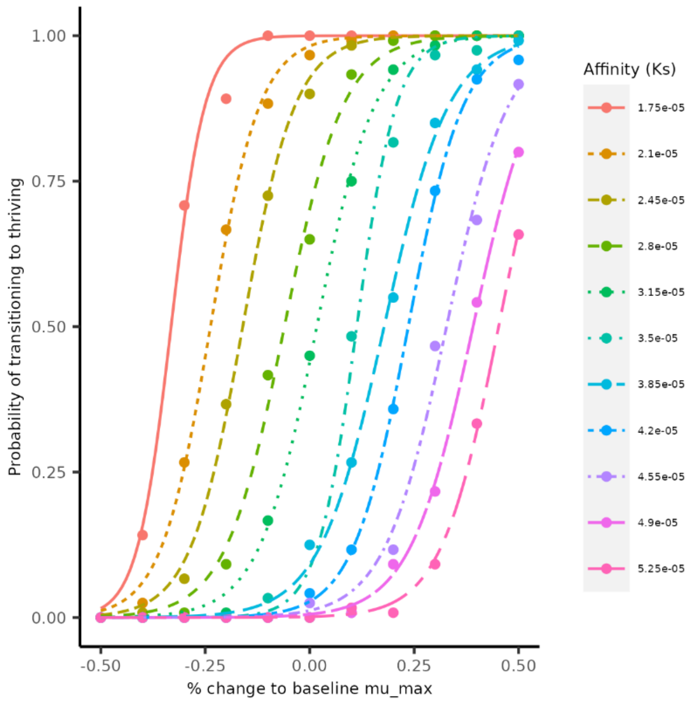


**Figure S4:** Sigmoid fits for 4 initial organisms spaced 10 diameters apart.

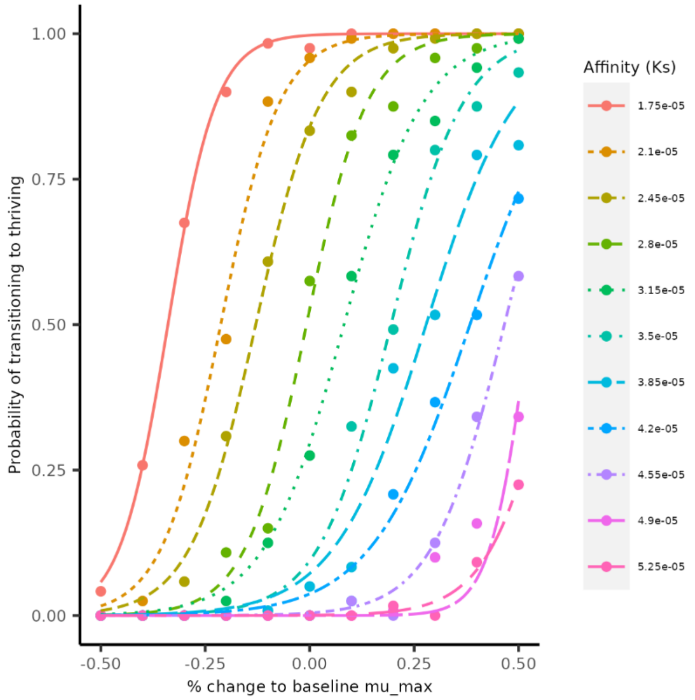


**Figure S5:** Sigmoid fits for 9 initial organisms spaced 2.5 diameters apart.

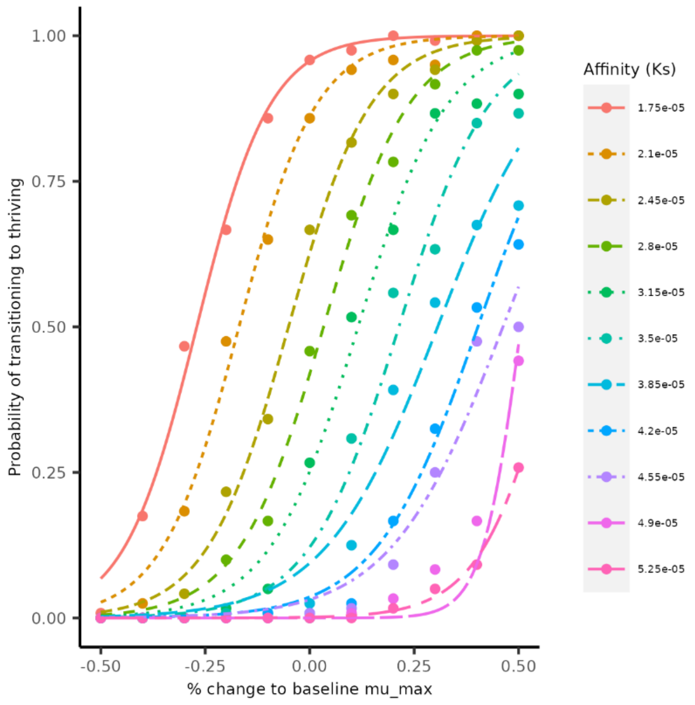


**Figure S6:** Sigmoid fits for 9 initial organisms spaced 5 diameters apart.

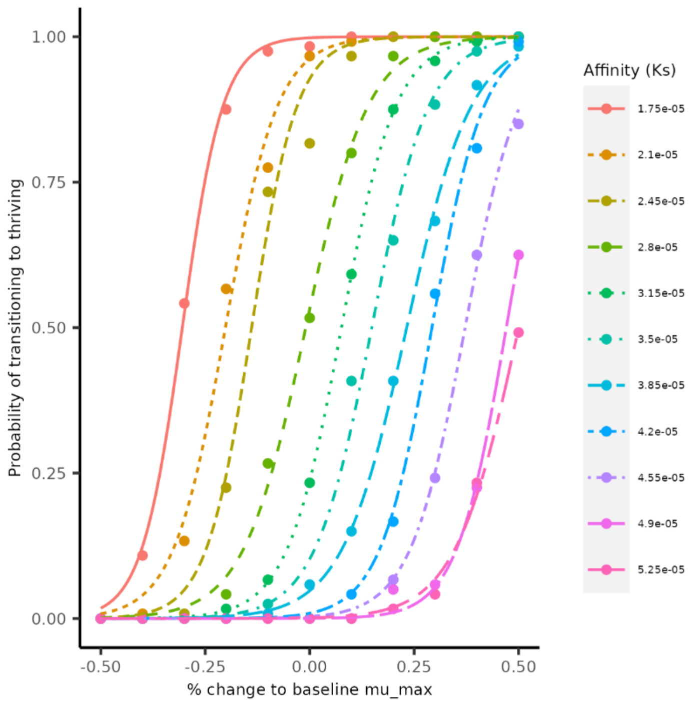


**Figure S7:** Sigmoid fits for 9 initial organisms spaced 10 diameters apart.

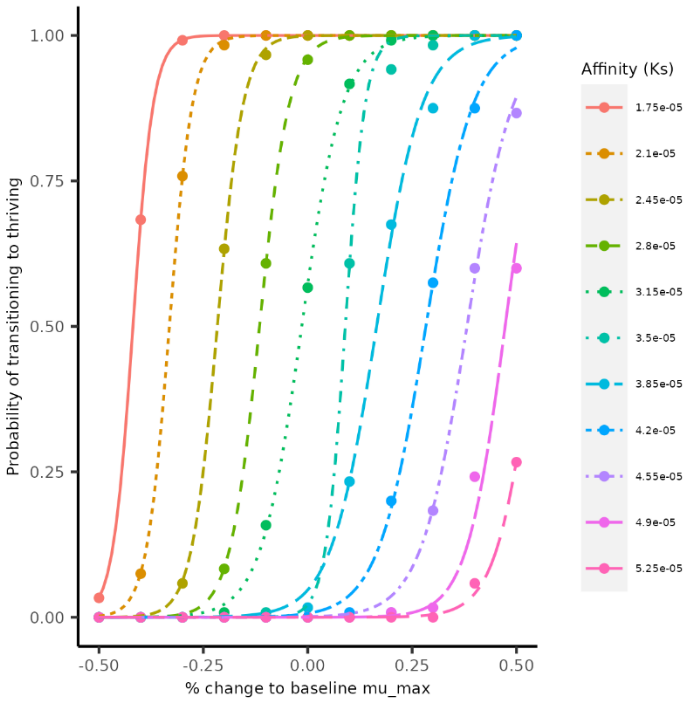


**Figure S8:** Sigmoid fits for 16 initial organisms spaced 2.5 diameters apart.

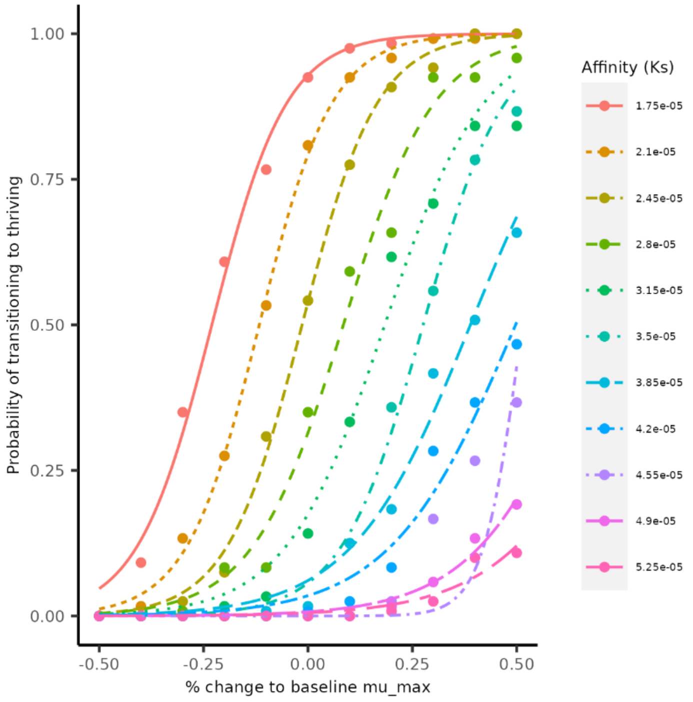


**Figure S9:** Sigmoid fits for 16 initial organisms spaced 5 diameters apart.

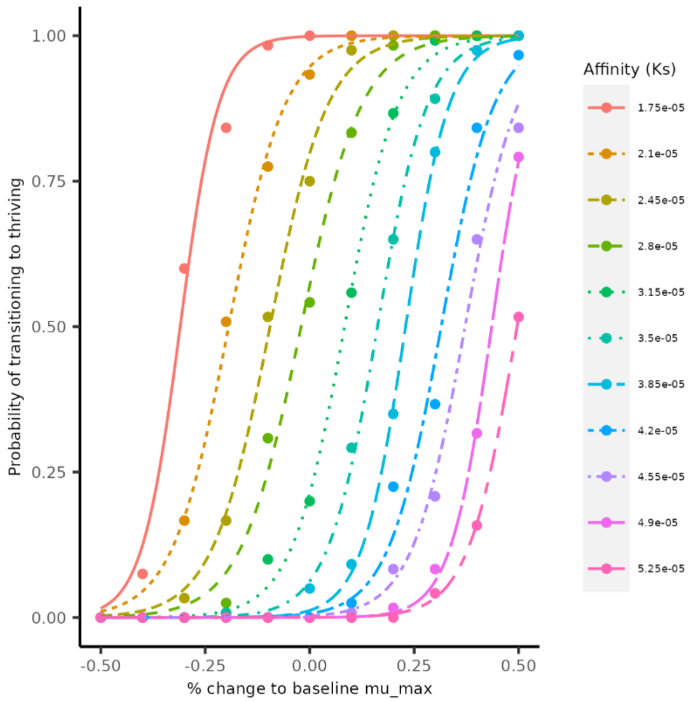


**Figure S10:** Sigmoid fits for 16 initial organisms spaced 10 diameters apart.

#### Model Selection and Fits

**Table S2:** Multiple Linear Regression Model candidates and fit metrics. Selected model indicated by *

| Model Description | Formula (R-Style) | Adjusted $R^2$ |
|---|---|---|
| Main effects only | `loglike ~ mu_pct + ks_pct + nbugs + spacing` | 0.82 |
| With 3-way interactions | `loglike ~ mu_pct + ks_pct + nbugs + spacing + mu_pct * ks_pct + mu_pct * nbugs + mu_pct * spacing + ks_pct * nbugs + ks_pct * spacing + nbugs * spacing + mu_pct * ks_pct * nbugs + mu_pct * ks_pct * spacing + mu_pct * ks_pct * nbugs * spacing` | 0.83 |
| Backwards step reduction from 3-way interactions* | `loglike ~ mu_pct + ks_pct + nbugs + spacing + mu_pct * spacing + ks_pct * spacing` | 0.82 |

**Table S3:** Coefficients and significance for selected MLR model. Bold indicates significance ($p < 0.05$)

| Coeffient (R-style) | Estimate | p-value |
|---|---|---|
| `(Intercept)` | -2.22684 | **3.25e-10** |
| `mu_pct` | 19.32634 | **< 2e-16** |
| `ks_pct` | -18.83760 | **< 2e-16** |
| `nbugs` | -0.07192 | **0.00362** |
| `Spacing` | -0.02581 | 0.50750 |
| `mu_pct:spacing` | 0.34685 | **0.00493** |
| `ks_pct:spacing` | 0.34402 | **0.00529** |

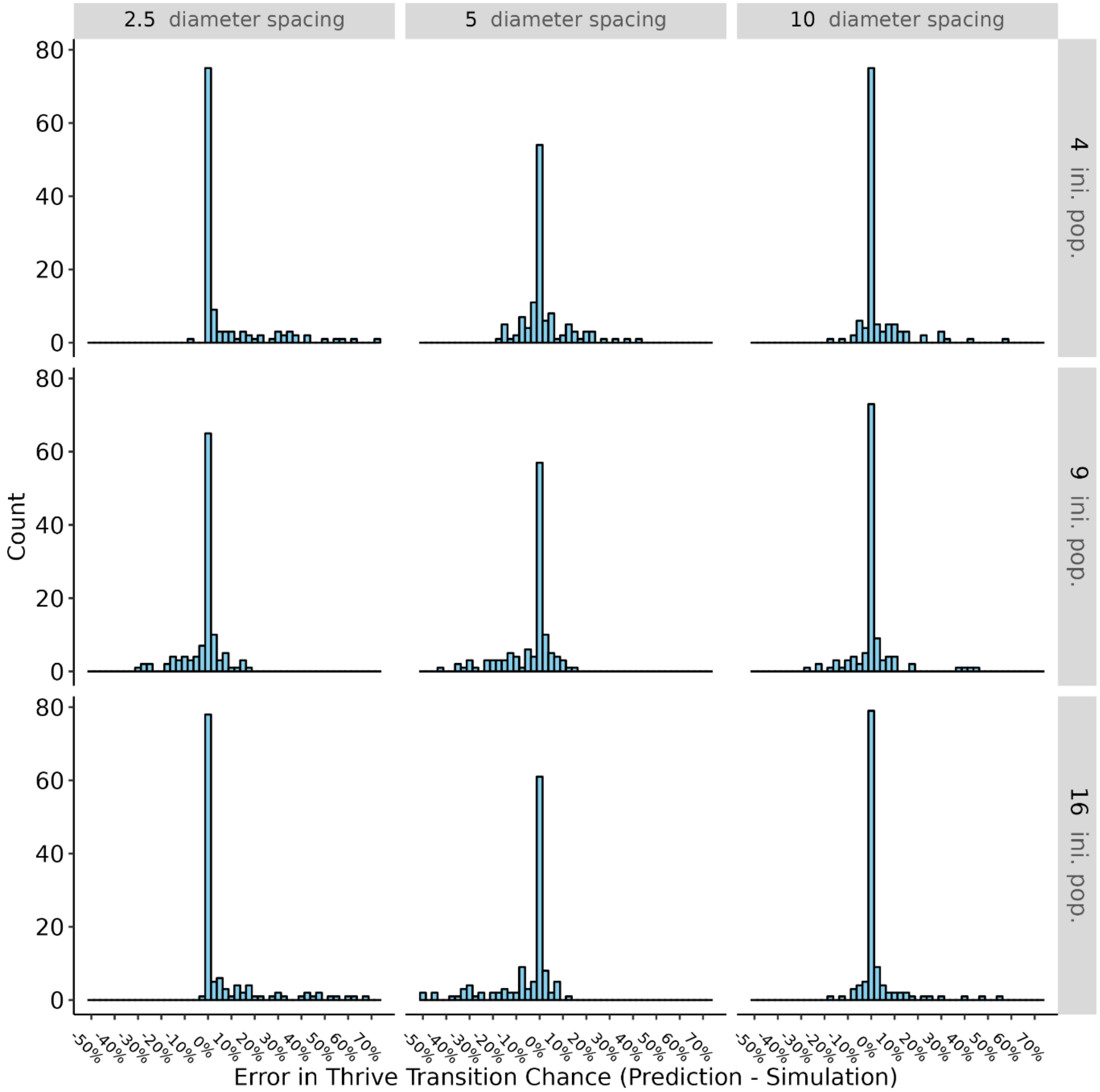


**Figure S11:** MLR prediction errors for each crowding condition

**Table S4:** Generalized Additive Model candidates and fit metrics. Selected model indicated by *.

| Model Description | Formula (R-style) | Adjusted $R^2$ | AIC |
|---|---|---|---|
| Main effects only, all smooths | `prob_thrive ~ s(mu_pct) + s(ks_pct) + s(nbugs, k = 3) + s(spacing, k = 3)` | 0.783 | -424 |
| Main effects only, all linear | `prob_thrive ~ mu_pct + ks_pct + nbugs + spacing` | 0.768 | -362 |
| Main effects only, crowding terms linear | `prob_thrive ~ s(mu_pct) + s(ks_pct) + nbugs + spacing` | 0.775 | -387 |
| With 2-way interactions, all smooth | `prob_thrive ~ s(mu_pct) + s(ks_pct) + s(nbugs, k = 3) + s(mu_pct, ks_pct) + s(mu_pct, nbugs) + s(ks_pct, nbugs)` | 0.956 | -654 |
| With 2-way interactions, crowding terms linear | `prob_thrive ~ s(mu_pct) + s(ks_pct) + nbugs + spacing + s(mu_pct, ks_pct) + s(mu_pct, by = nbugs) + s(mu_pct, by = spacing) + s(ks_pct, by = nbugs) + s(ks_pct, by = spacing) + nbugs * spacing` | 0.947 | -1923 |
| With 3-way interactions, all smooth | `prob_thrive ~ s(mu_pct) + s(ks_pct) + s(nbugs, k = 3) + s(spacing, k = 3) + s(mu_pct, ks_pct) + s(mu_pct, nbugs) + s(mu_pct, spacing) + s(ks_pct, nbugs)+ s(ks_pct, spacing) + s(nbugs, by = spacing, k = 3) + te(mu_pct, ks_pct, spacing, d = c(1, 2)) + te(mu_pct, ks_pct, nbugs, d = c(1, 2))` | 0.978 | -2797 |
| With 3-way interactions, crowding terms linear | `prob_thrive ~ s(mu_pct) + s(ks_pct) + nbugs + spacing + s(mu_pct, ks_pct, k = 60) + s(mu_pct, by = nbugs) + s(mu_pct, by = spacing) + s(ks_pct, by = nbugs) + s(ks_pct, by = spacing) + nbugs * spacing + te(mu_pct, ks_pct, by = spacing) + te(mu_pct, ks_pct, by = nbugs)` | 0.953 | -2017 |
| Backwards step reduction from 3-way interactions, all smooth* | `prob_thrive ~ s(mu_pct) + s(ks_pct) + s(nbugs, k = 3) + s(spacing, k = 3) + s(mu_pct, ks_pct, k = 60) + s(mu_pct, spacing) + s(ks_pct, spacing) + s(nbugs, by = spacing, k = 3) + te(mu_pct, ks_pct, spacing, d = c(1, 2)) + te(mu_pct, ks_pct, nbugs, d = c(1, 2))` | 0.979 | -2850 |

**Table S5:** Term information and significance for selected GAM. Bold indicates significance ($p < 0.05$)

| Term (R-style) | k’ | Extended Degrees of Freedom | p-value |
|---|---|---|---|
| `s(mu_pct)` | 9 | 1.000 | 0.12326 |
| `s(ks_pct)` | 9 | 1.000 | 0.65427 |
| `s(nbugs)` | 2 | 1.819 | 0.06761 . |
| `s(spacing)` | 2 | 1.000 | **1.64e-06** |
| `s(mu_pct,ks_pct)` | 57 | 46.141 | **< 2e-16** |
| `s(mu_pct,spacing)` | 21 | 4.713 | **0.00029** |
| `s(ks_pct,spacing)` | 21 | 7.112 | **< 2e-16** |
| `s(nbugs):spacing` | 3 | 2.979 | **< 2e-16** |
| `te(mu_pct,ks_pct,spacing)` | 89 | 48.812 | **< 2e-16** |
| `te(mu_pct,ks_pct,nbugs)` | 97 | 45.657 | **< 2e-16** |

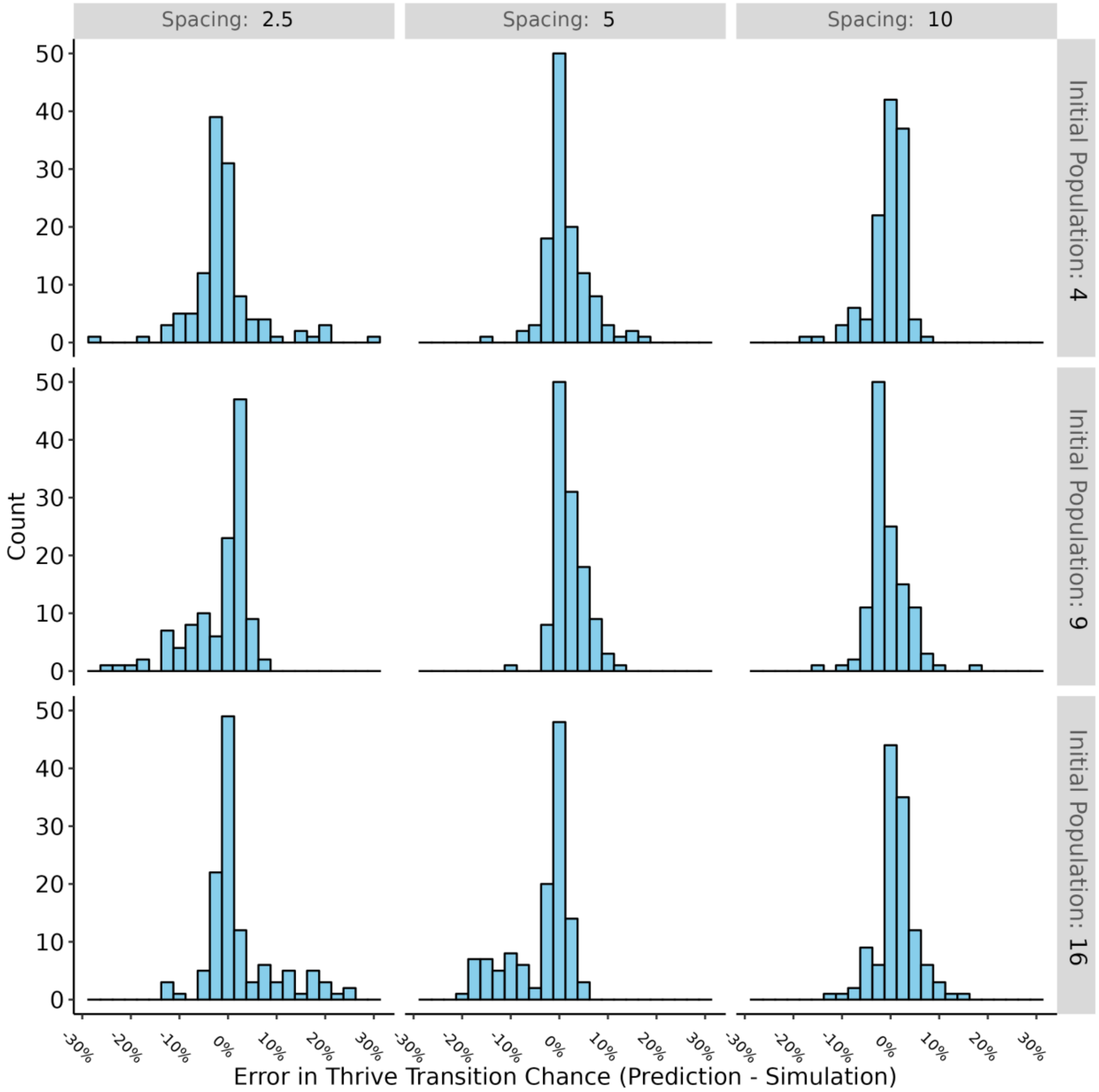


**Figure 12:** GAM prediction error distributions by crowding condition

#### Test of Underlying Assumptions

**Table S6:** Chi-squared test p-values for counts of 'biggest loser' lineages at each initial site location. A significant p-value ($\alpha_{corrected}$=0.0056) would suggest that a particular initial site location may be biased towards failure.

| Initial Population | Spacing (Diameters) 2.5 | 5 | 10 |
|---|---|---|---|
| 4 | 0.036 | 0.52 | 0.90 |
| 9 | 0.047 | 0.83 | 0.76 |
| 16 | 0.72 | 0.98 | 0.26 |

### Additional Parameter Trend Analysis

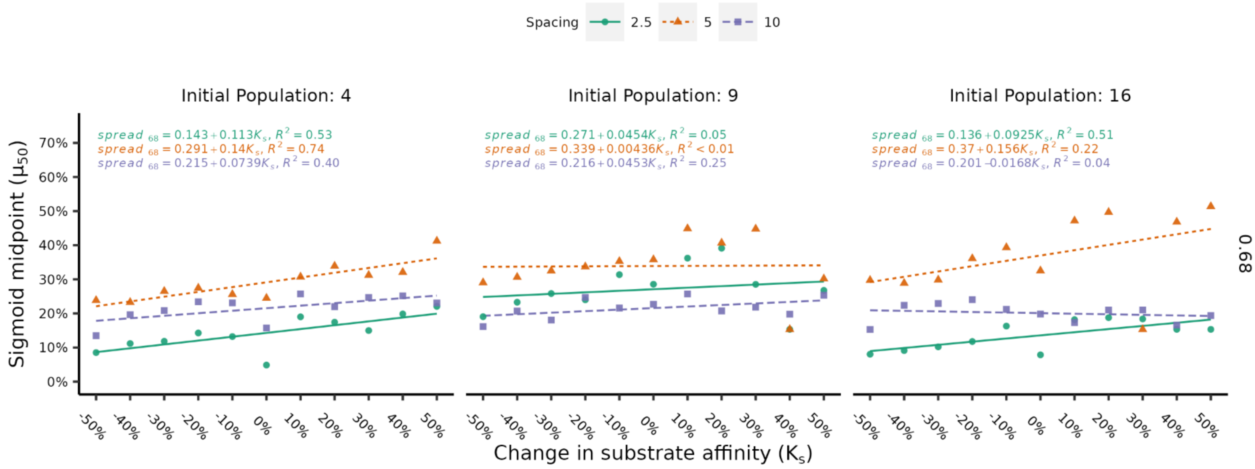


**Figure S13:** Under each crowding condition, *spread68* changes with *KS*. Insofar as trends are present, moderate spacing produces the widest *spread95* and the differences between spacings increases with population size.

### Fitted Model Larger-Resolution Images

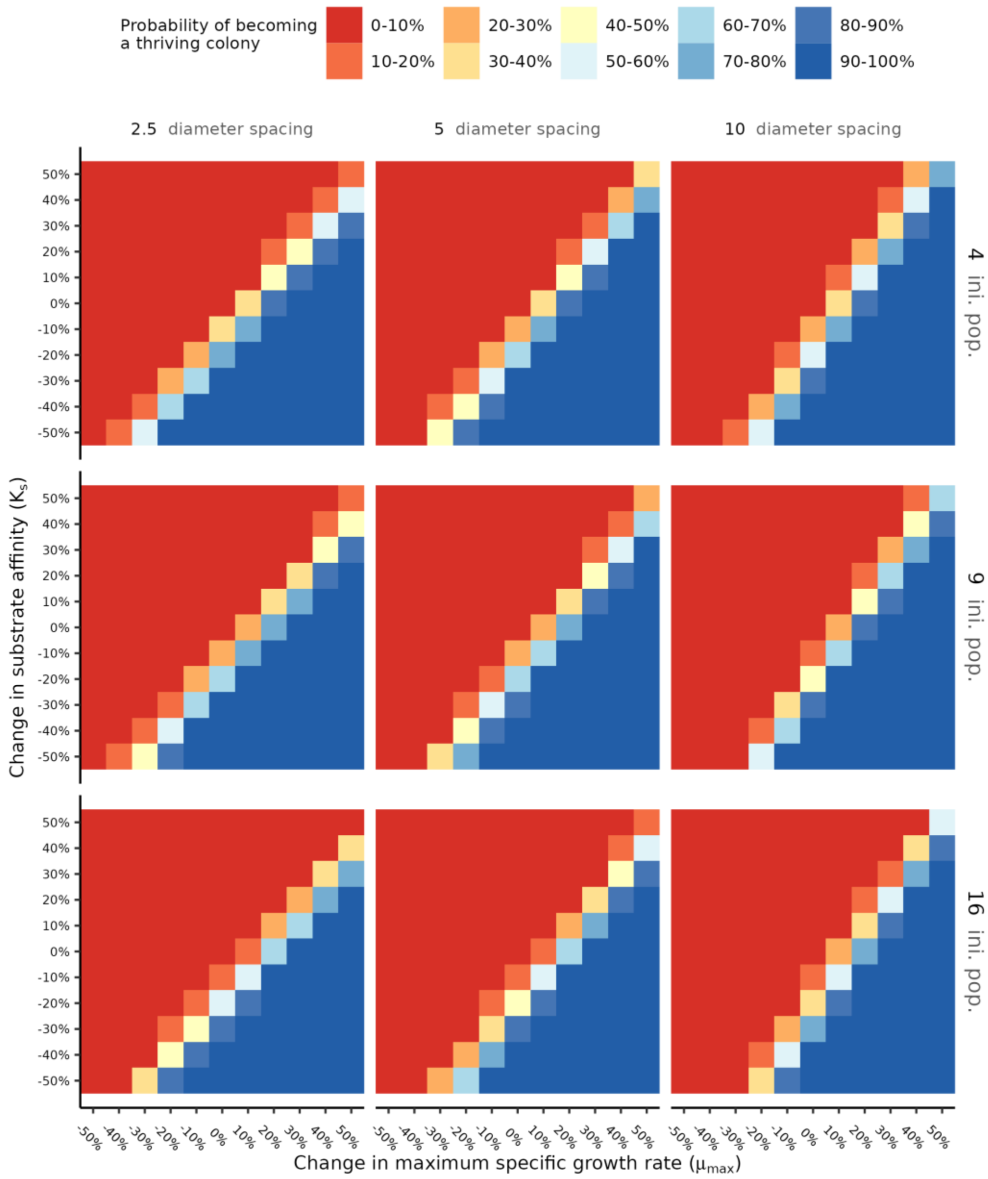


**Figure S14:** MLR model predictions, large format of Figure 7, panel A

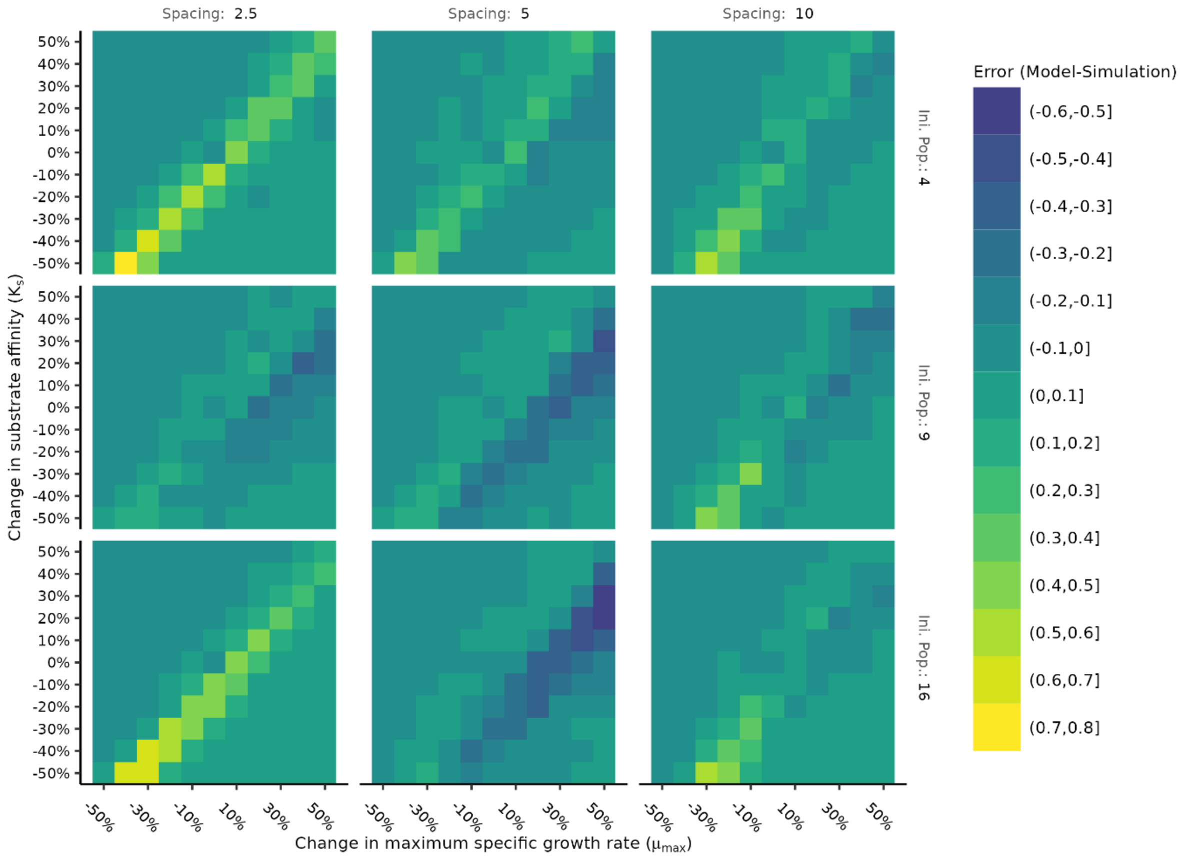


**Figure S15:** MLR model error. Large format version of Figure 7, panel D

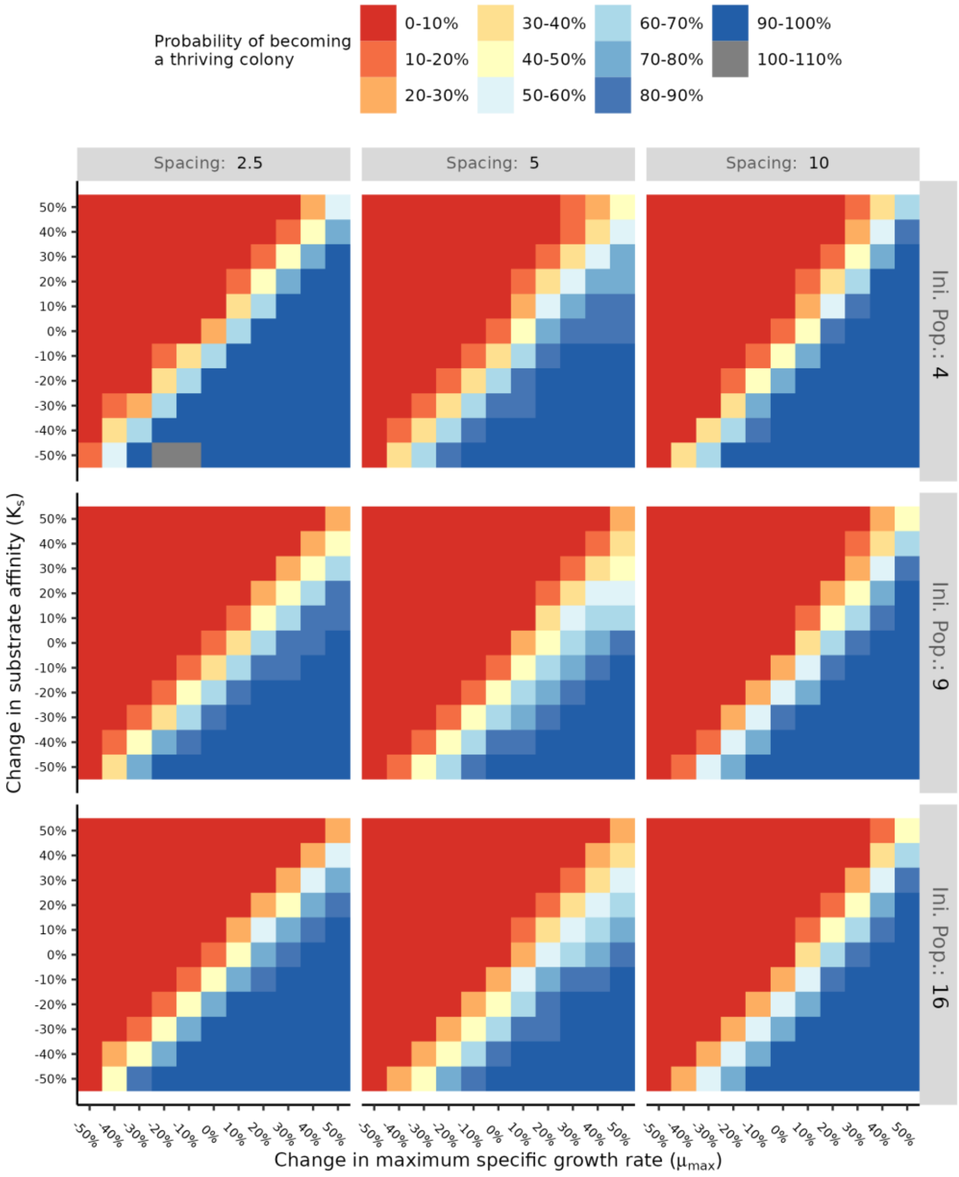


**Figure S16:** GAM predictions, large format of Figure 7, panel B.

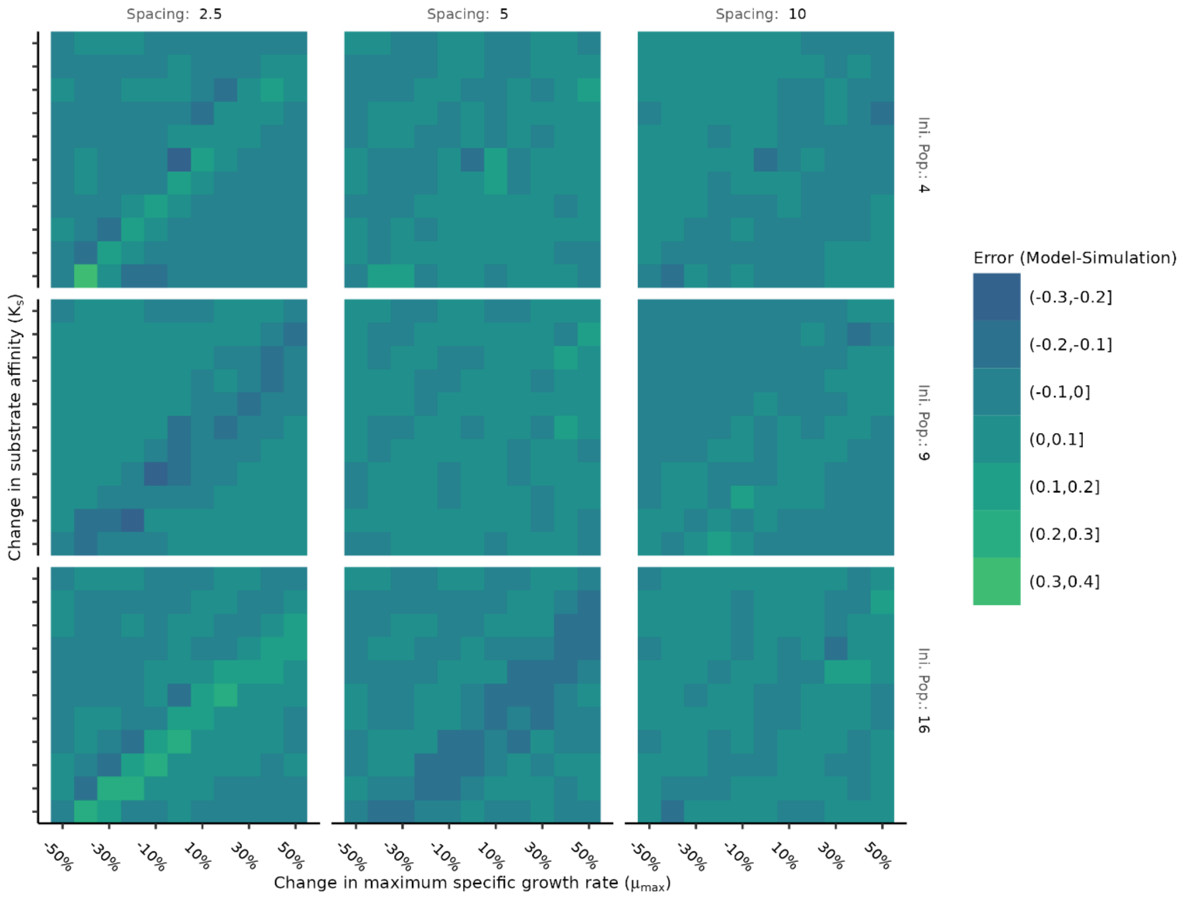


**Figure S17:** GAM error. Large format version of Figure 7, panel E
